# Multiple testing correction over contrasts for brain imaging

**DOI:** 10.1101/775106

**Authors:** Bianca A. V. Alberton, Thomas E. Nichols, Humberto R. Gamba, Anderson M. Winkler

## Abstract

The multiple testing problem arises not only when there are many voxels or vertices in an image representation of the brain, but also when multiple contrasts of parameter estimates (that is, hypotheses) are tested in the same general linear model. Here we argue that a correction for this multiplicity must be performed to avoid excess of false positives. Various methods have been proposed in the literature, but few have been applied to brain imaging. Here we discuss and compare different methods to make such correction in different scenarios, showing that one classical and well known method is invalid, and argue that permutation is the best option to perform such correction due to its exactness and flexibility to handle a variety of common imaging situations.

## 1. Introduction

A well known problem in brain imaging is the multiplicity of tests, which arises given the fact that a statistical test is performed in each voxel or vertex of an image representation of the brain. However, an equally common situation in which such multiplicity occurs is when multiple contrasts of parameter estimates of the same general linear model (GLM) are considered, or even many multiple different such models. In effect, data acquisition is one of the most expensive and laborious stages of an experiment, such that often the same data are reused and reanalysed in different ways to consider different models and hypotheses. If left uncontrolled, such multiplicity can lead to an undesirably high number false positives.

For example, in the Human Connectome Project (HCP), the task fMRI N–back working memory experiment is conducted using different classes of stimuli (faces, places, tools and body parts) (Barch et al., 2013). Analyses involving these classes can be corrected using Bonferroni, but results would be overly conservative due to dependence among the tests. Comparisons among the classes using methods such as analysis of variance (ANOVA) followed by pairwise comparisons do not control the error rate, as we demonstrate later in this paper.

Even though a number of methods have been proposed for correction for similar problems in non-imaging fields (Hochberg and Tamhane, 1987; Hsu, 1996), most of these have seen little use in the brain imaging. In this short technical note, we assert that such correction is necessary, discuss and compare a few existing methods through which it can be implemented, and provide an approach based on permutation tests that provides an exact (as opposed to conservative) control over the error rate, even when the multiple hypotheses being tested are not independent.

## 2. Theory

### 2.1. Notation and general aspects

Consider the general linear model (GLM) expressed by:

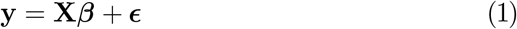

where **y** is *N* × 1 vector containing the image data for *N* subjects at a given voxel (or vertex), **X** is the *N* × *R* design matrix with the modeled *R* explanatory variables, ***β*** is the *R* × 1 vector with the (to be estimated) parameters that scale the variables in **X** so as to explain the variability observed in **y**, and *ϵ* is the *N* × 1 vector of random errors. The coefficients can be estimated via ordinary least squares as 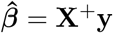, where the symbol ^+^ represents the Moore–Penrose pseudo-inverse. The interest is to test *K*, *K* ≥ 1, null hypotheses, each of them represented as 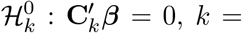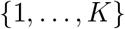, where **C**_*k*_ is a *R* × *S*_*k*_ full rank matrix that defines the contrasts of parameter estimates. In ANOVA designs, the interest is a global (*omnibus*) null hypothesis of no difference among all groups, which can be tested using the *F*-statistic:

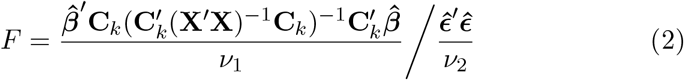

where *ν*_1_ = rank(**C**_*k*_) and *ν*_2_ = *N* − rank(**X**) are the degrees of freedom. If 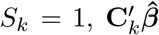 is scalar and a *t*-statistic can be obtained from *F* as 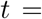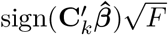, or equivalently:

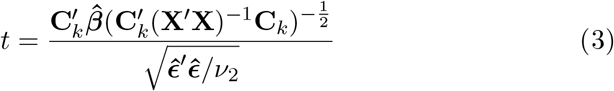

The test is said to be *significant* at the level *α*, 0 ≤ *α* ≤ 1, if the probability of observing a random variable *T* larger or equal than *t* (or *F*), if the null hypothesis is true, is smaller or equal than *α*, i.e., ℙ(*T* ≥ *t*) ≤ *α*. This is the p-value, and can be computed based on distributional assumptions on *t*, or through a resampling procedure, such as a permutation test (Pesarin and Salmaso, 2010).

Permutation is a non-parametric method in which the distribution of the test statistic is ascertained by explicitly calculating all (or a large number of) the possible values that it could assume should the null hypothesis be true. These values are obtained by randomly rearranging the data as if the null hypothesis were indeed true (Nichols and Holmes, 2002; Winkler et al., 2014). For a number *J* of permutations of the data, each one associated with a test statistic *t*_*j*_, *j* = {1, 2, …, *J*}, the p-value can be calculated as the number of occurrences of a random (after permutation) *t*_*j*_ that is larger or equal to the observed, original statistic *t* (obtained without any permutation) divided by the number of permutations performed. In other words, *t*_*j*_ takes the role of the random variable *T* described above. Note that *t* itself (computed from the model without any permutation) should also be counted among the set of *J* statistics computed after permutation, such that the smallest possible p-value in a permutation test cannot be smaller than 1/*J*, and therefore cannot be zero (Phipson and Smyth, 2010). Permutation tests are particularly interesting in experiments in which there are only a few subjects, or if the assumptions that underlie parametric tests cannot be confirmed (Ludbrook and Dudley, 1998).

### 2.2. Multiple testing

As more tests are conducted, the more likely it will be that at least one will be declared significant even if no actual effect exists. The problems associated with the multiplicity of tests across points (e.g., voxels or vertices) in an image representation of the brain are well known, and various strategies have been devised (for reviews, see Nichols and Hayasaka, 2003; Farcomeni, 2008; Nichols, 2012). In general, approaches target the control of one of two different error quantities: the *family-wise error rate* (FWER), which is defined as the chance of any false positive across all tests, and the *false discovery rate* (FDR), which is the expected proportion of false positives across all tests in which an effect has been found significant.

When only one test is considered (that is, in the absence of multiple testing), all that is needed for a decision on rejecting or not the null hypothesis are the p-value *p* and a (usually pre-defined) test level *α*. When more than one test is performed, either the test level can be *corrected* (*α*_cor_), in which it is changed so as to accommodate the multiplicity of tests (the p-values remain unchanged), or the p-values can be *adjusted* (*p*_adj_), in which these are changed instead (the test level remains then unchanged). In either case, the modifications are such that the FWER or the FDR is controlled at the level *α* (though for FDR, test levels are often denoted as *q*).

Multiple image points, such as voxels or vertices, are not the only way in which multiple testing can occur in brain imaging; various other sources of multiplicity that are not simply multiple spatial tests are possible; some of us have previously used the term *multiple testing problem type II* (MTP-II), so as to distinguish them from the usual multiplicity due to the many measurements taken across space (MTP-I)^l^ (Winkler et al., 2016b). The multiplicity of contrasts of model parameter estimates belongs to that class. As with correction over voxels or vertices (MTP-I), which is performed across all image points of interest (e.g., the whole brain, or within a region of interest), it is desirable that the correction in the MTP-II considers only the hypotheses of interest. In other words, it is not always the case that all and every possible contrast of parameter estimates is relevant or meaningful to be tested; for example, when testing *G* groups, one may only be interested in each group individually (*G* tests), or the interest may lie on differences among specific groups, and not necessarily on all the *G*(*G* − 1) possible pairwise group comparisons.

Methods to account for the multiplicity of contrasts include the same that can be used over any set of p-values, such as Bonferroni or FDR, or can be specific to the context of the general linear model. Below we briefly summarise some of these methods (see also Table 1). Although here the focus is on FWER, the conceptualization of the correction across contrasts, as well as the application of permutation tests, remains similar for FDR.

**Table 1:** Summary comparison of methods for correction across contrasts.

| Method | Global test p-value | Local tests' adjusted p-values |
| --- | --- | --- |
| Dunn–Šidák | — | $1 - (1 - p_{\text{unc}})^\tau$ |
| Bonferroni | — | $\tau \cdot p_{\text{unc}}$ |
| Fisher's LSD | $1 - F_{\text{cdf}}(F; G - 1, \nu_2)$ | $1 - t_{\text{cdf}}(t; \nu_2)$ |
| Tukey | — | $1 - Q_{\text{cdf}}(t\sqrt{2}; G, \nu_2)$ |
| Scheffé | — | $1 - F_{\text{cdf}}\left(\frac{\text{sign}(t) t^2}{G-1}; G - 1, \nu_2\right)$ |
| Fisher–Hayter | $1 - F_{\text{cdf}}(F; G - 1, \nu_2)$ | $1 - Q_{\text{cdf}}(t\sqrt{2}; G - 1, \nu_2)$ |
| Westfall–Young | — | $1 - t_{\text{cdf}}^{\max}(t)$ |
| Wang–Cui | $1 - F_{\text{cdf}}^{\text{WC}}(F; G - 1, \mathbf{n})$ | $1 - Q_{\text{cdf}}(t\sqrt{2}; G - 1, \nu_2)$ |

Most of these methods are performed in a single, local step that produces the final, adjusted p-value; others require an initial global (*omnibus*) test, which, if significant, is followed by local tests (also known, in this context, as *post hoc* tests). If the initial global test is not significant, then the second step the p-value is not computed, and can be treated as $p_{\text{adj}} = 1$ , that is, not significant. Fisher's LSD and Fisher–Hayter require an initial $F$ -test with $(G - 1)$ and $\nu_2$ degrees of freedom. The global test in Wang–Cui use the $F$ -statistic compared against a null distribution computed through Monte Carlo methods, and which depends on the number of groups and also on a vector $\mathbf{n}$ that contains the sizes of each group (thus explicitly accommodating unbalancedness). Note that the adjustment in Bonferroni can lead to p-values larger than 1, which reflects the fact that this method is an approximation.

#### Dunn–Šidák

The probability that two independent events can occur simultaneously is given by the product of their probabilities. Thus, when multiple independent statistical tests are conducted, the corrected test level can be computed as *α*_cor_ = 1 − (1 − *α*)^1/*/τ*^, where *τ* is the number of tests (when the correction is applied to multiple contrasts, and using our notation, *τ* = *K*). This idea was considered by Tippett (1931), and later proved in a related context by Dunn (1958) and Šidák (1967). The Dunn–Šidák (DS) equation is often described as an adjustment of one or more p-values instead of correcting *α*, that is, *p*_adj_ = 1 − (1 − *p*_unc_)^*τ*^, where *p*_unc_ is the original (uncorrected) p-value obtained in a given test. In this case, the FWER is controlled for the adjusted p-values without the need to modify the test level *α*. The equality holds only if the tests are independent; the adjustment is conservative otherwise.

#### Fisher’s least significant difference

This is a two-step procedure suggested by Fisher (1935) as a way to identify which tests were responsible for driving the overall (*omnibus*) result of an ANOVA (it does not appear that Fisher was concerned with multiple testing when proposing this test). In this method, an *omnibus F*-test is performed to detect if there are any group differences; if and only if the *F*-test is significant at *α*, a second stage is performed, in which follow up (*post hoc*) *t*-tests between each pair of groups are evaluated at the same level *α*. However, it has been demonstrated (Hayter, 1986) that the maximum probability of finding at least one incorrect result can greatly exceed the test level, growing rapidly as the number of groups exceed 3, and is therefore not recommended (Hsu, 1996). The test remains valid for up to and including 3 groups, though.

#### Bonferroni

Based on Boole’s inequality, which states that, given a series of events, the probability that at least one happens is smaller or equal to the sum of the probabilities of each of the individual events, Bonferroni (1936) suggested an approximation for the corrected test level as *α*_cor_ = *α*/*τ*. The simplicity and intuitive appeal of this approximation were certainly determinants to its enormous popularity across all scientific domains over many decades. However, even for independent tests, Bonferroni’s method is slightly conservative when compared to Dunn–Šidák (which is exact). If the tests are not independent, the correction becomes yet more conservative. For the relation between Bonferroni and Dunn–Šidák, see Appendix A.

#### Tukey

For one-way ANOVA layouts, Tukey (1953) proposed that p-values for the comparison between each pair of group means could be computed with reference to the studentized range distribution (SRD), that is, 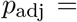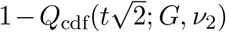, where *Q*_cdf_ represents the cumulative distribution function (cdf) from the SRD, *G* is the number of groups and *ν*_2_ is the number of degrees of freedom. Unlike Fisher’s LSD, this method does not require an initial *F*-test. This procedure assumes that all possible pairwise comparisons could be of interest, where the greatest difference between means is the most likely to be rejected. When the groups are unbalanced, the procedure is known to be conservative, and the degree of conservativeness varies with the number of groups and the severity of the unbalance (Hochberg and Tamhane, 1987). Although this method has been applied mainly for ANOVA designs, it could also be considered for scenarios in which multiple comparisons are performed among regression coefficients for continuous variables (e.g., comparing different continuous signals against a single reference regressor). This method, and variants of it, have received a number of different names in different settings, including *T-Procedure*, *wholly significant difference* (WSD), *honestly significant difference* (HSD), and, when applied to unbalanced models, *Tukey–Kramer test*. For simplicity, here we call it simply “Tukey” for both balanced and unbalanced experiments.

#### Scheffé

Under the null hypothesis that all group means are equal, the adjusted p-value can be obtained as 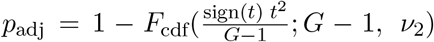, Where *F*_cdf_ is the cumulative distribution function of the *F* distribution, with *ν*_1_ = *G* − 1 and *ν*_2_ degrees of freedom, and sign(*t*) is the sign of *t* statistic (*F*_cdf_ evaluates negative values as zero). This method was proposed by Scheffé (1953) as a way to correct the statistic across all possible contrasts. Therefore, its correction is much more stringent than the Fisher’s LSD and Tukey’s method, being very conservative when pairwise comparisons are performed, but has the advantage of correcting across contrasts modelling more complex comparisons among groups.

#### Fisher–Hayter

Hayter (1986) proposed that the p-value could be adjusted by changing the second step of Fisher’s LSD to accommodate the SRD as 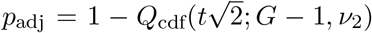. In comparison with Tukey’s method, this procedure has greater power due to the loosening of *G* to *G* − 1, and still maintains control of the FWER due to the *F*-test applied in the first step of the procedure. Without this first step, the FWER would be slightly greater than the defined *α* level. Being a modification of Fisher’s LSD, this method is sometimes named Fisher–Hayter (FH) procedure (Seaman et al., 1991).

#### Westfall–Young (permutation)

Using permutation tests to calculate the distribution of the maximum statistic across the set of tests that are being corrected, the adjusted p-value can be computed as 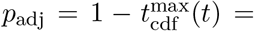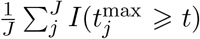, where *J* is the total number of permutations, *I*(*·*) is the indicator function and 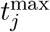 is the maximum value of *t*_*j*_ across the contrasts^2^ (Westfall and Young, 1993). The reason why this works is that the test whose statistic is maximum across all tests being considered is also the one that has the maximum statistic for any subset of these tests that includes it, such that this approach constitutes a *closed testing procedure* (Marcus et al., 1976). Such closed procedures are known to strongly control the FWER.

#### Wang–Cui

It has been suggested that a lower critical level for the first stage (the *F*-test) of the Fisher–Hayter method can be found in a single, holistic procedure that integrates the results of all possible pairwise group comparisons from the second stage. To accomplish this, a Monte Carlo distribution of the *F*-statistic is computed under the assumption of complete normality but, crucially, for a given Monte Carlo realization *m*, whenever 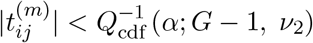 for all *i* ≠ *j* groups, *F*^(*m*)^ is defined as 0, as opposed to the actual calculated *F*-statistic for that realization. Wang and Cui (2017) argue that this procedure is more powerful than Fisher–Hayter, being exact for balanced designs, and conservative otherwise (yet more powerful than Fisher–Hayter).

### 2.3. Use in permutation tests

In principle, all methods discussed above and summarized in Table 1 could be used with permutation tests. Bonferroni and Dunn–Šidák, which use as starting point the uncorrected p-values, can be adjusted regardless of whether the p-values are obtained parametrically or non-parametrically; for these methods, all that is needed is independence among the tests and that, under the complete null hypotheses, the p-values are uniformly distributed; if tests are not independent, the resulting inferences are conservative, yet valid. Fisher’s LSD, Tukey, Scheffé, Fisher–Hayter and Wang–Cui, as originally proposed, require that the test statistic at one or two stages are compared to some reference distribution, under the assumption of normality. However, nothing prevents that these distributions are also obtained non-parametrically, via permutations. For example, the *F*-distribution used in Fisher’s LSD, Fisher–Hayter, and Scheffé can be obtained applying a permutation test using all possible unique (linearly independent) comparisons among the groups. For Tukey, Fisher–Hayter and Wang–Cui, the SRD *Q*-distribution can likewise be obtained using the the permutation distribution of the largest difference among all comparisons, using only permutations that change subjects among those groups that are being tested in a given contrast (Petrondas and Gabriel, 1983). For each of these cases, a specific algorithm would be built, and permutations could make these tests robust to departures from normality.

The Westfall–Young method stands distinct from these other, potential methods in that it is intrinsically non-parametric, whereby the distribution of the maximum statistic is obtained via permutations. As with such other potential methods, normality is not assumed, nor independence, since the synchronized permutations over which the maximum statistic is obtained implicitly captures eventual dependencies among the tests. Further, the procedure is optimal in that it targets the very definition of familywise error rate: if the maximum is significant, then surely at least one rejection of the null hypothesis has happened; if the null is true for all tests, then that is a familywise error. The distribution of the maximum statistic is, therefore, a direct way to control the FWER. Moreover, it is algorithmically simpler than permutation versions of various of the other tests, and requires the permutation data only for the hypotheses of interest. The Westfall–Young is the *de facto* method for permutation-based FWER-corrected inference. Henceforth, and consistent with the literature, when referring to correction using permutations, we are referring to this method.

## 3. Evaluation Methods

### 3.1. Synthetic data

To investigate the error rates and power of each method, we considered one-way ANOVA designs, with simulated data representing 2000 voxels (these could also be construed as vertices, or any other type of imaging element). The dataset consisted of random variables following a normal distribution with zero mean and unit variance. We considered 2500 realisations of 8 different scenarios where three simulation parameters were manipulated: presence or absence of signal, balanced or unbalanced designs, and the correction over all possible pairwise group comparisons or only the largest subset of linearly independent contrasts.

With respect to signal, when it was added, it was to all voxels of subjects in groups 1 and 2, and with size defined as **X***β* with *β* = [*β*_1_, *β*_2_, 0, …, 0]^′^, where *β*_1_ = +*s*_*t*_, *β*_2_ = −*s*_*t*_, and 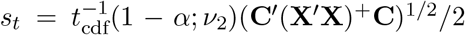. **C**= [+1, −1, 0, …, 0]^*′*^ is the contrast vector and *α* = 0.05 is the test level. By adding a positive signal to group 1 and a negative signal to group 2, both with magnitudes determined by *s*_*t*_, an approximate power of 50% irrespective to the sample size or number of groups can be expected in all simulations for the contrast comparing these two groups before any adjustment has been made for the multiplicity of tests (in this case, multiplicity of contrasts). Details on how to add signal that confers a specific power before the correction are described in Appendix B. Also, although stronger effects are expected to be detected in the comparisons between groups 1 and 2, smaller effects can also be detected when groups 1 or 2 are compared with any other group. Thus, power calculations below consider effects in any comparison involving groups 1 or 2. The contrasts used to calculate power were excluded from the FWER calculation.

The simulations used sample sizes of 40, 50, 70 or 90 subjects. For the balanced scenarios, subjects were divided in 4, 5, 7 or 9 groups respectively, always with 10 subjects per group. For the unbalanced scenarios, subjects were likewise divided into 4, 5, 7 or 9 groups, for the same respective sample sizes, however with each group consisting of a random number of participants under the constraint that each group had at least 4 subjects.

The comparisons considered (1) all possible pairwise group differences and (2) a subset of linearly independent comparisons. In the latter case, we tested the hypotheses that group 3 were greater than every one of the other groups. The reason to investigate a correction over a subset of independent contrasts is that all possible pairwise group differences imply dependencies among them (for example, for three groups, the difference between groups 2 and 3 is fully determined once differences between groups 1 and 2, and 1 and 3 are known). Such dependencies do not exist for a subset of contrasts that are independent from each other, and by selecting all contrasts involving group 3, we obtain the minimum set of contrasts that can be tested while representing all possible groups. This type of model is very common, for example, when testing various patient groups against one control group, and is also a case that demonstrates that not all possible pairwise group comparisons might be of interest. For the analyses using this subset, power can be calculated as the proportion of detected true effects in contrasts where group 3 is greater than groups 1 and 2. The greatest proportion of signal is expected in the contrast testing that group 3 is greater than 2, which gives an approximate (expected) power of 20.6%. The FWER is estimated as the proportion of tests in which the null hypothesis is rejected for contrasts that compare whether group 3 is greater than each of the groups from 4 to *G*.

All methods were evaluated using custom code written in GNU Octave (Eaton et al., 2014). Tukey, Fisher–Hayter and Wang–Cui correction, however, used critical values pre–calculated with the functions “ptukey” and “qtukey” from the R statistical software (R Core Team, 2018), whereas Westfall–Young permutation method used PALM – Permutation Analysis of Linear Models (Winkler et al., 2014) with 5000 permutations. Bonferroni and Dunn–Šidák were applied over parametric p-values computed using Student’s *t* distribution.

### 3.2. Real data

We used data from the Healthy Brain Network (HBN) (Alexander et al., 2017) made publicly available by the Child Mind Institute (CMI, New York, NY, USA) to evaluate how correction over the number of contrasts could affect a realistic data analysis. The data collection, as well their distribution in anonymized format, was approved by their institutional ethics review board. Structural, *T*_1_-weighted magnetic resonance images were processed using FreeSurfer (Dale et al., 1999; Fischl et al., 1999). Quality control was performed based on visual inspection of the reconstructed pial and inflated surfaces, with particular emphasis on the subjects with extreme values for the metrics produced by the tool MRIQC (Esteban et al., 2017), and most extreme Euler numbers for either of the hemispheres (Rosen et al., 2018); the threshold for the Euler number was −80, and subjects in which either hemisphere had values more negative than this threshold were excluded. This allowed selection of 278 subjects that successfully finished the FreeSurfer processing, that passed quality control, and further, that had complete data for the variables that we selected to be used in the statistical analysis (described below).

As dependent variables we investigated the volume of the subcortical structures automatically segmented by FreeSurfer: thalamus, caudate, putamen, pallidum, hippocampus, amygdala, nucleus accumbens, and ventral diencephalon (VDC, a group of structures whose precise limits are not typically discernible with *T*_1_-weighted scans, and that includes hypothalamus, mammillary bodies, lateral and medial geniculate nuclei, subthalamic nuclei, substantia nigra and nucleus ruber) (Fischl et al., 2002, 2004). We also investigated associations with the estimated total intracranial volume (eTIV) (Buckner et al., 2004).

As independent variables, we chose two disparate measures for investigation, one related to social factors (Barratt simplified measure of social status, BSMSS) (Barratt, 2006) and another related to physiological factors (extra-cellular water, ECW). Two analyses of covariance (ANCOVA) designs were considered, one to investigate BSMSS and another for ECW. The subjects were divided in 5 groups using the 20th, 40th, 60th and 80th percentiles of each of these two variables, so that the grouping of subjects of BSMSS was different from the grouping of ECW. By dividing the continuous variable into discrete units, we can more easily consider an AN(C)OVA scenario for which some of the correction methods were originally devised (even though no such limitation exists for Dunn–Šidák, Bonferroni, Scheffé or permutation tests), and further, we can accommodate the possibility of certain non-linear effects. It should be emphasised, however, that this division is completely arbitrary, and is done here solely for convenience. Age (5 – 15 years, mean = 9.85, standard deviation = 2.65) and sex (180 males and 98 females) were included as nuisance variables. All pairwise differences were tested. Contrasts 1 through 4 tested whether group 1 would be larger than the others, contrasts 5 through 8 tested whether group 2 would be larger than the others, and so forth, for a total of 20 contrasts covering all possible pairwise group comparisons in both directions (Figure 1).

**Figure 1:**
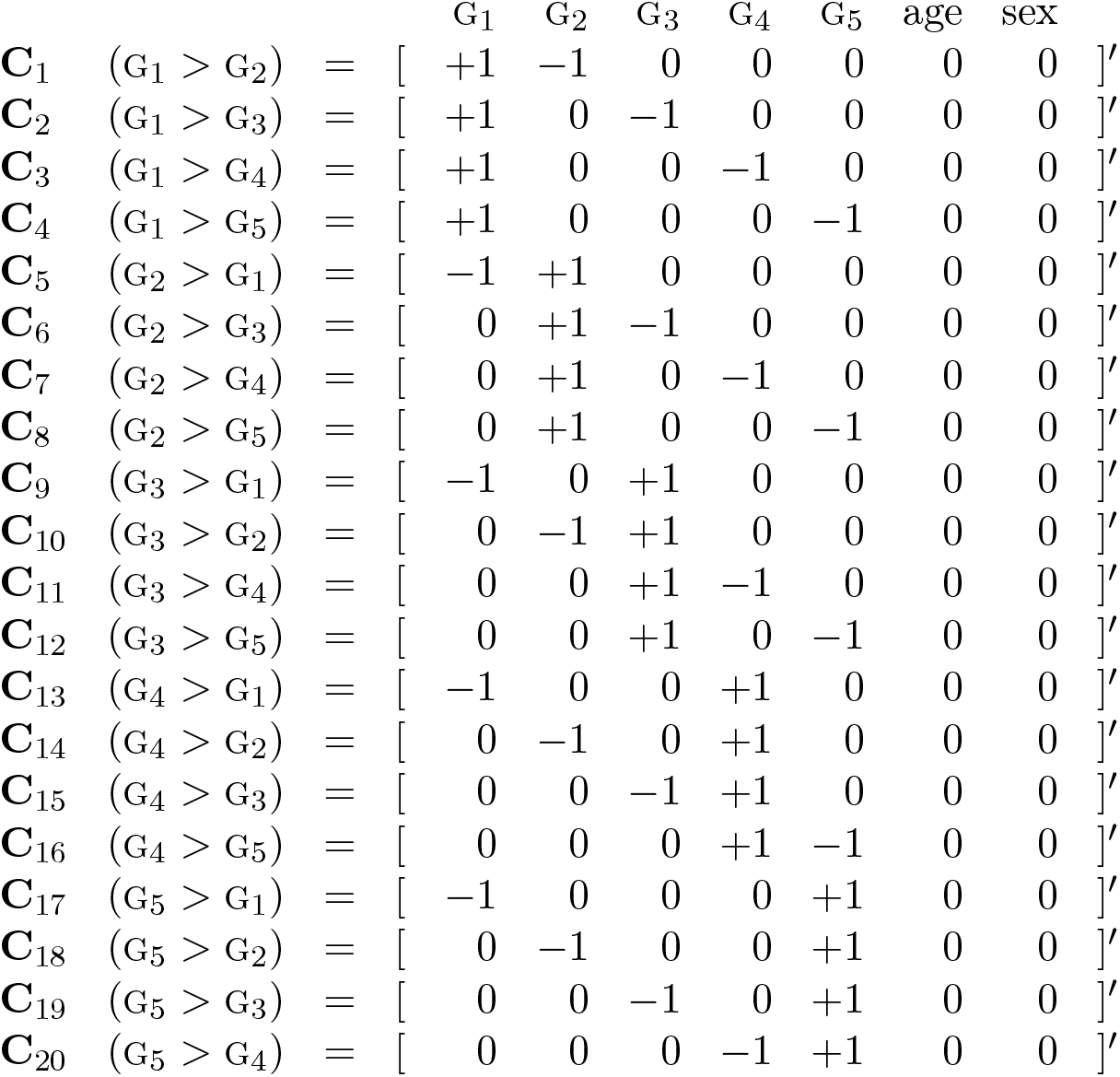
List of contrasts testing all possible pairwise comparisons among the 5 groups (G_1_, …, G_5_) as encoded by the design matrix (not shown). The last two regression coefficients modelled age and sex, in an ANCOVA design.

Statistical analysis used PALM with 10000 permutations and tail approximation (Winkler et al., 2016a) to calculate p-values, both uncorrected (for contrasts) and corrected with Westfall–Young permutation method. All other methods were applied using custom code written on GNU Octave. Bonferroni and Dunn–Šidák were applied over parametric p-values using Student’s *t* distribution. As with the synthetic data, for Tukey, Fisher–Hayter and Wang–Cui we invoked the functions “ptukey” and “qtukey” from R.

## 4. Results

### 4.1. Synthetic data

Figure 2 shows the error rates and power of each simulation testing all pairwise comparisons. All evaluated procedures controlled the FWER in the absence of signal, that is, when the null hypothesis was true for all *K* = *G*(*G* − 1) contrasts, for both balanced and unbalanced models.

**Figure 2:**
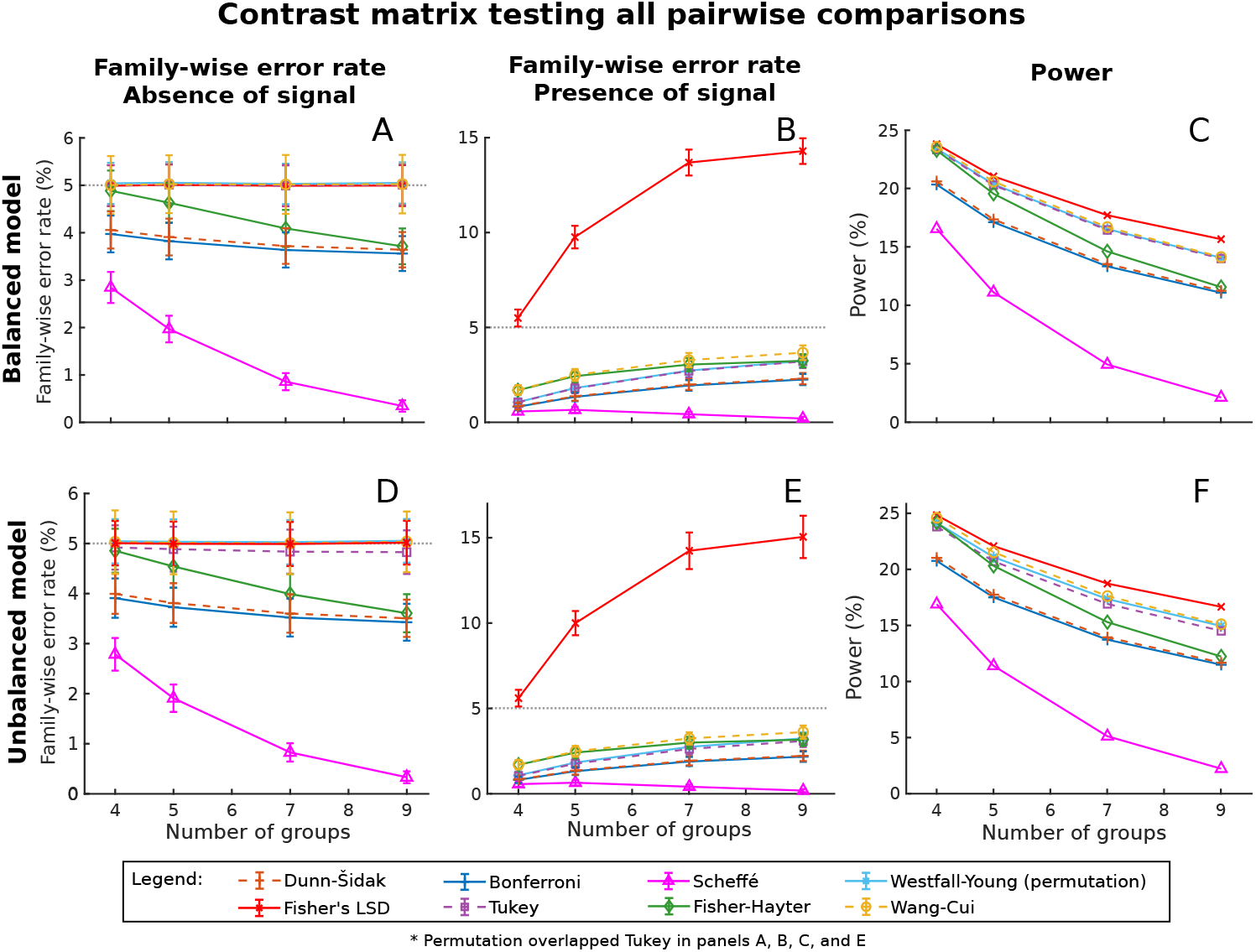
Mean family-wise error rate and power after correcting across contrasts using Dunn–Šidák, Fisher LSD, Bonferroni, Tukey, Scheffé, Fisher–Hayter, Westfall–Young permutation method and Wang–Cui when testing all pairwise comparisons. Starting with balanced models, (A) shows the FWER results in the absence of signal when all contrasts are considered, (B) shows the FWER in the presence of signal, but considering the contrasts that have no signal, and (C) the respective power, i.e., the ability to detect signal for the contrasts that had signal; panels (D), (E), and (F) show, respectively, the same, for unbalanced models. The error bars in panels A, B, C and E represent the standard deviation of the mean error rate. A power of 50% was expected before any correction was performed.

However, in the presence of signal for some of the contrasts, LSD substantially exceeded the error rate of 5% for those contrasts that did not have signal (Figure 2, B and E). Although with 4 groups the FWER is only slightly above the nominal level (5.51% and 5.62% for balanced and unbalanced groups, respectively), with 9 groups the FWER reaches 14.30% for balanced and 15.07% for unbalanced models, which is around three times higher than the expected nominal level of 5%. All other methods maintained the control over the FWER in the presence of signal for some of the contrasts. Permutation and Tukey had a similar FWER for all group configurations, as well as Fisher–Hayter and Wang–Cui when 4 and 5 groups were used. Bonferroni had a FWER slightly smaller than Dunn–Šidák, with a ratio around 0.9770 between them^3^. The most conservative method was Scheffé, with the highest observed FWER of 0.67%, when 5 groups were considered.

When all possible pairwise comparisons were analysed, and considering only the methods that control the FWER at the nominal level, Wang–Cui had the greatest power, with of 23.58% and 24.61% for balanced and unbalanced models, respectively, when 4 groups were considered. Permutation had a slightly smaller power of 23.41% and 24.16% for balanced and unbalanced models, and was closely followed by Tukey, which had an observed power of 23.28% and 23.82% for balanced and unbalanced models. Fisher–Hayter started with a competitive power of 24.18% in unbalanced models with 4 groups, but as the number of groups increased, the power of Fisher–Hayter decreased at a faster rate than the other methods (Figure 2, C and F). Dunn– Šidák, with a power of 21.04% when 4 unbalanced groups were considered, showed a slightly greater power than Bonferroni, with a power of 20.76% in the same configuration. Scheffé was the most conservative method, with a power of 16.56% in balanced and 16.91% unbalanced models with 4 groups.

In general, the FWER and the power had similar values for experiments with both balanced and unbalanced designs. The exception was in the simulation with complete absence of signal (Figure 2, D), in which Tukey exhibited a small reduction in the observed FWER and power that was proportional to the increase of number of groups in the unbalanced model. However, the same trend did not appear in the FWER in the presence of signal.

The greatest difference in the performance of the various methods appeared when only a subset of linearly independent contrasts was used (Figure 3). In this case, Tukey, Fisher–Hayter and Wang–Cui were very conservative and had a power smaller than 1.4% in the simulation with 8 groups. In the subset of linearly independent contrasts, Westfall–Young had the greatest power among the valid methods, 18.85% and 11.35% in the experiment with 4 groups, for balanced and unbalanced respectively.

**Figure 3:**
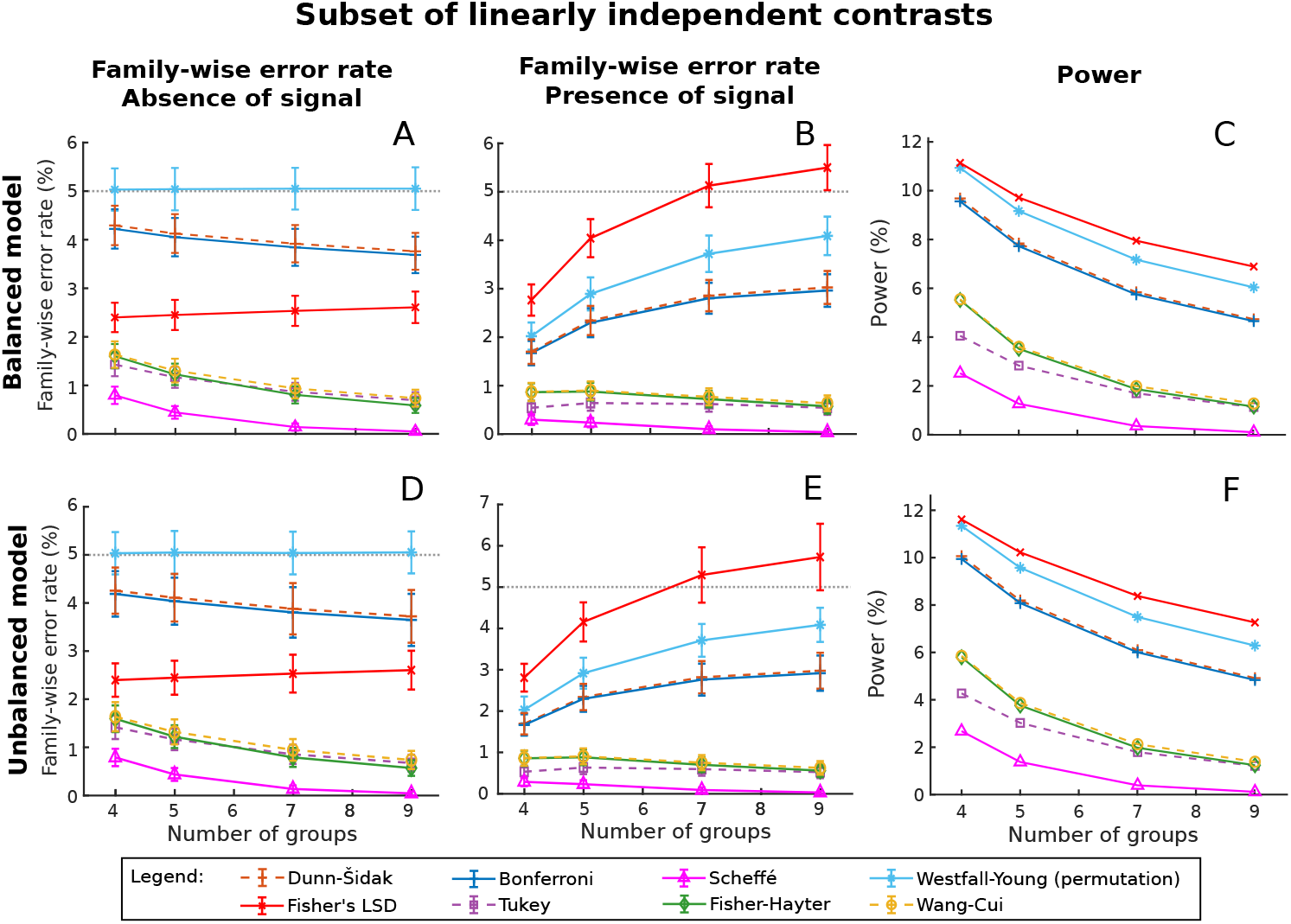
Mean family-wise error rate and power after correcting across contrasts using Dunn–Šidák, Fisher LSD, Bonferroni, Tukey, Scheffé, Fisher–Hayter, Westfall–Young permutation method and Wang–Cui when testing only a subset of linearly independent contrasts. Starting with balanced models, (A) shows the FWER results in the absence of signal when all contrasts are considered, (B) shows the FWER in the presence of signal, but considering the contrasts that have no signal, and (C) the respective power, i.e., the ability to detect signal for the contrasts that had signal; panels (D), (E), and (F) show, respectively, the same, for unbalanced models. The error bars in panels A, B, D and E represent the standard deviation of the mean error rate. A power of 20.6% was expected before any correction was performed.

### 4.2 Real data

Division of the subjects into 5 groups using the 20th, 40th, 60th and 80th percentiles resulted in unbalanced groups with 56 subjects on average. Table 2 shows the range from BSMSS and ECW, as well as the values used to divide the subjects into these discrete groups.

**Table 2:** Measures and its percentiles used to divided the data into groups.

| Measure | Range | Percentiles |  |  |  |
| --- | --- | --- | --- | --- | --- |
|  |  | 20th | 40th | 60th | 80th |
| BSMSS | 9 – 66 | 35.05 | 48 | 54.5 | 61 |
| ECW (liters) | 4.23 – 49.59 | 11.28 | 15.39 | 19.24 | 26.20 |

Without correction for the multiplicity of contrasts, some effects were detected among BSMSS groups (Figure 4): subjects in group 2 had greater mean cortical volume in the left nucleus accumbens than the subjects from groups 3 and 4 (contrasts 6 and 7); group 4 showed greater mean volume in the right pallidum than group 1 (contrast 14); and subjects in group 5 had greater estimated total intracranial volume, as well as larger thalamus, hippocampus, amygdala, pallidum and VDC volumes than some of the other groups (contrast 16 to 20). After correction, no such effects were observed.

**Figure 4:**
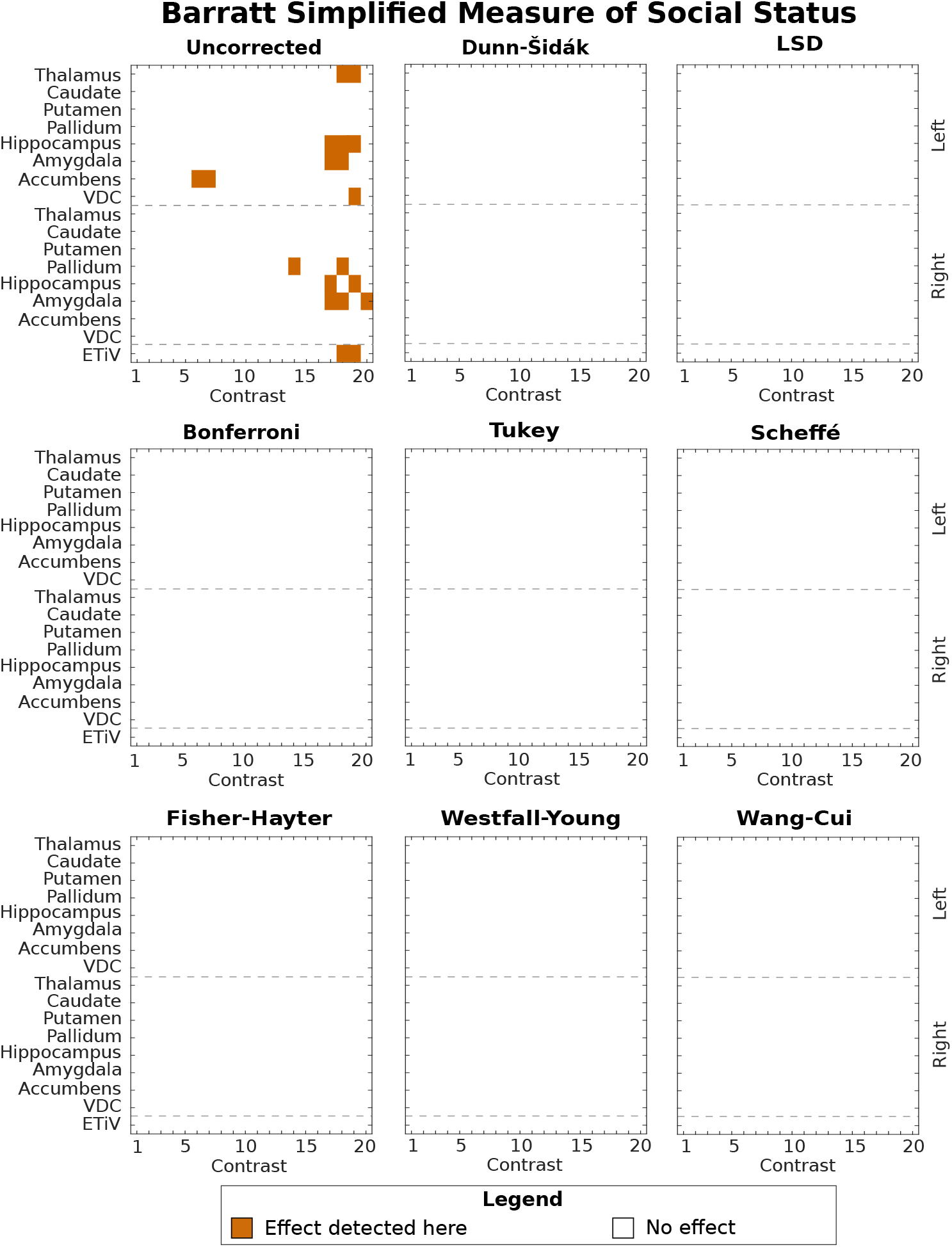
Effect in both hemispheres per contrast detected when dividing the subjects into five groups using the percentiles from BSMSS.

For ECW, a number of regions were found to have significantly larger volumes for all ECW groups in relation to group 1 (contrasts 5, 9, 13 and 17). From these, eTIV, as well as right pallidum, amygdala and accumbens did not survive the correction across contrasts. These results are summarised in Figure 5. Among the correction methods, Fisher–Hayter and Wang–Cui led to the same results. The correction performed with Westfall–Young and Tukey had similar performance to each other, differing only in the detection of greater left amygdala volume in group 2 in relation to group 1 (contrast 5). Permutation results differed a bit more when compared with those obtained with Fisher–Hayter, in which a difference in the volume of the left accumbens that was significant with Westfall–Young was no longer so after Fisher–Hayter (contrast 9, G_3_ > G_1_), and another effect was significant with Fisher–Hayter in the right caudate, but not with permutation (also contrast 9). A much sparser set of results was observed with the Dunn–Šidák, Bonferroni and Scheffé approaches. Dunn–Šidák did not identify the same effects as permutation in the left hemisphere in the amygdala, hippocampus, ventral diencephalon and caudate (contrasts 5, 9 and 17), nor the same effects that remained after the correction with Fisher–Hayter in the right caudate. Further, Bonferroni did not detect the effect in the pallidum (contrast 17). Scheffé detected the smaller amount of effects, namely in the thalamus, left pallidum, and right hippocampus (contrasts 9 and 13). While a substantial number of comparisons remained significant after Fisher’s LSD correction, the simulations had already demonstrated that these results are invalid; nonetheless, these are also shown in Figure 5.

**Figure 5:**
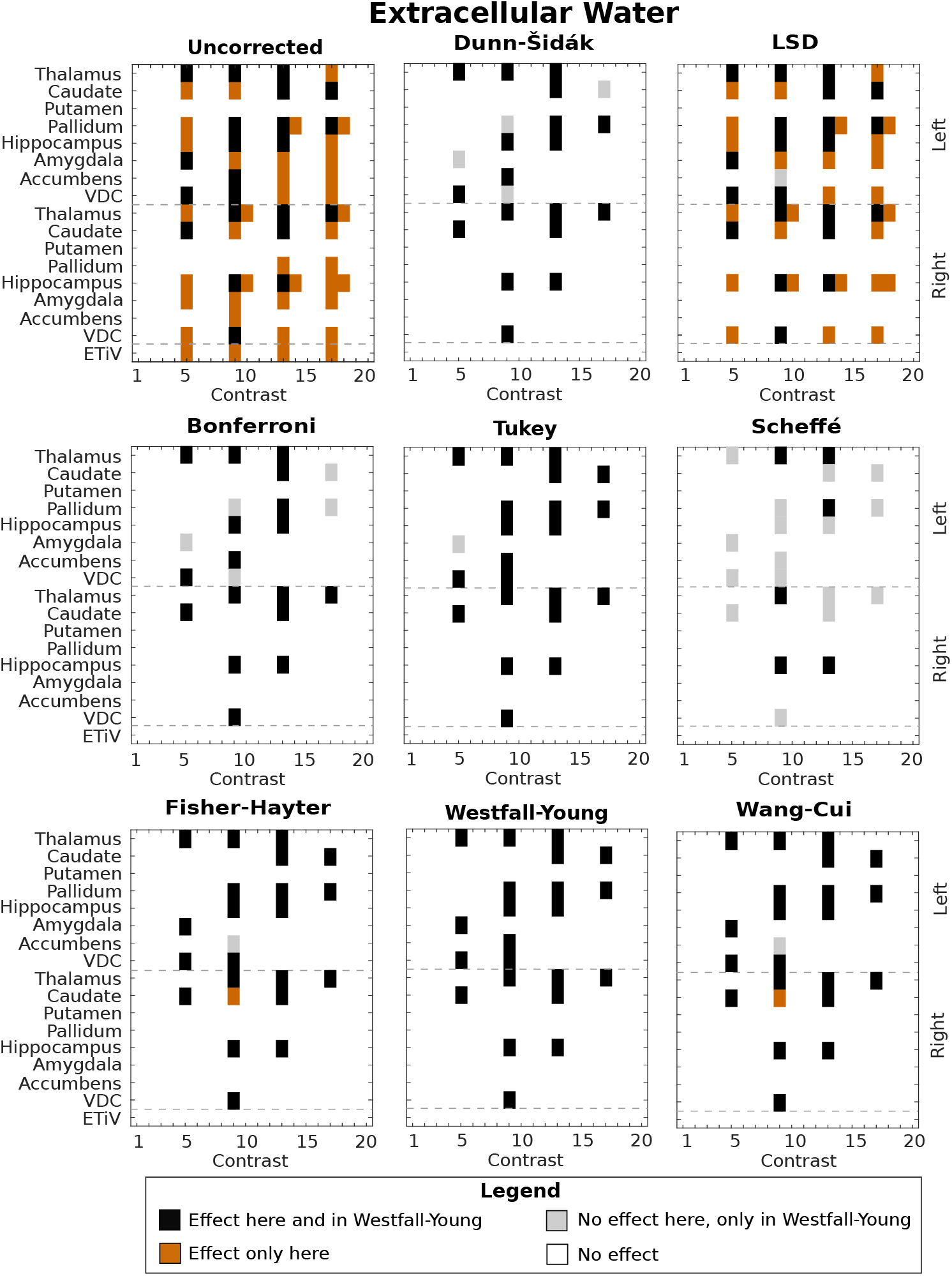
Effect in both hemispheres per contrast detected when dividing the subjects into five groups using the percentiles from extracellular water.

## 5. Discussion

### 5.1. Error rates and power

The Westfall–Young permutation method was the only that performed well in all simulation scenarios: it always controlled the FWER at the test level (was an exact procedure). While methods such as Wang–Cui, Fisher– Hayter and Tukey had a good performance in some scenarios, they have limitations: they do not extend trivially to designs that do not follow an AN(C)OVA design, they take into account all possible pairwise comparisons and do not perform as well when only a subset of them are of interest. Moreover, Fisher–Hayter and Wang–Cui require, along with LSD, an initial *F*-test, whereas for the others this step is bypassed altogether, with the correction applicable directly to the *t*-tests.

When some of the pairwise comparisons contained signal, LSD vastly exceeded the 5% error rate for the comparisons that did not contain signal, confirming the results obtained by Hayter (1986), and showing that LSD offers only weak control of the FWER, i.e., it controls the FWER only when no true effect is present in any of the hypotheses being tested, and becoming invalid (that is, extrapolating the test level) when signal is present in some of the contrasts. The other methods assessed, in turn, offered strong control when their assumptions were met, that is, they ensured an FWER equal to or smaller than the test level, both in the absence and presence of true effects (for the definition of strong and weak control, see Hochberg and Tamhane, 1987; Nichols and Hayasaka, 2003).

Among the methods tested, Scheffé led to the lowest observed FWER among all methods in all scenarios. When all pairwise group comparisons are considered, that is, the case in which there are dependencies among the tests, Dunn–Šidák and Bonferroni were also very conservative, being substantially below the nominal test level of 5%, resulting in low power. With linearly independent contrasts, however, Dunn–Šidák and Bonferroni became less conservative, and their difference when compared to permutation was always below 1.5%. Between Dunn–Šidák and Bonferroni, the former is more powerful, and is exact when the tests performed are independent (whereas even in these cases, Bonferroni is slighly conservative, not exact).

In the complete absence of signal, as the number of groups increased, the FWER for most methods either remained stable, or tended to become more conservative, except Fisher’s LSD. With signal in groups 1 and 2, the observed FWER tended to approach the test level as the number of groups increased, presumably due to the smaller proportion of contrasts that contain signal. While this is not surprising, the rate with which these methods approached the test level differed substantially.

Although we focused on pairwise comparisons, AN(C)OVA designs may include more complex contrasts involving multiple groups, as for example, averaging two groups and comparing them against a third group. Contrasts may also test hypotheses that involve arbitrary combination of continuous and discrete regressors, or some contrasts comparing groups while others test continuous regressors (for example, in a design where categorical groups, age and sex are modelled, there may be interest in testing group differences, as well as linear and/or quadratic effects of age, or sex differences). Such heterogeneity, that goes beyond mere group comparisons, is not easily accommodated by methods based on the SRD, such as Tukey, Fisher–Hayter or Wang–Cui, but are handled easily by the Westfall–Young permutation method, which is very general.

From the simulation results presented in Figures 2 and 3, it is clear that, although there are differences in performance of the different methods, as more tests are conducted, the more strict the correction applied to the contrasts becomes. Although here the focus is on controlling the FWER, ensuring a high power, i.e., lower false negative rate, is equally important, and thus, a careful definition of the research hypotheses must precede statistical analysis. This, again, favors permutation methods, and in this case also Dunn–Šidák and Bonferroni. The reason is that all other other methods perform a broad correction that implicitly considers all possible pairwise comparisons (or even any possible comparisons, as in Scheffé), many of which might not be of any interest, and that unduly penalizes power (Figure 3, panels C and F).

### 5.2. Contrast correction and brain imaging

Accommodating correction over contrasts with brain imaging requires that both types of multiple testing, that is, across space (MTP-I), and across contrasts (MTP-II) (Winkler et al., 2016b) are considered together. For the former, permutation tests offer a solution that is valid, powerful, and with minimal assumptions (Westfall and Young, 1993; Nichols and Holmes, 2002; Winkler et al., 2014), and that extends to the latter in a quite simple manner: the correction can use the distribution of the maxima across imaging units (voxels, vertices, regions) *and* also across contrasts. This is simple enough to be included in any permutation testing algorithm.

The same cannot be said for the other methods discussed: corrections that would bypass the need for permutations for both contrasts and imaging units would need to rely not only on non-permutation methods for correction across contrasts for AN(C)OVA designs such as those presented here, but also on the many assumptions associated with methods as the random field theory (RFT; Worsley et al., 1996) for correction across image points. Furthermore, there are no known RFT results for fields following the SRD. The converse, that is, correcting first for the number of image points, and then the correction over contrasts, would find other difficulties since, likewise, there are no known results for the Euler characteristic across multiple, possibly non-independent, search volumes in the context of the RFT. All these would impose substantial challenges to guarantee control over the FWER. The most direct way to solve either of these is to use a permutation test, and once that is is used for one kind of multiple testing, correction for the other can be included in the same algorithm, with negligible further computational overhead.

### 5.3. Real data

An example of how contrast correction can be applied to an ANCOVA was shown, in this case after generating discrete groups by dividing subjects into subsets using the percentiles of two continuous variables. Although such correction should be done even when testing continuous regressors in the GLM, Fisher’s LSD, Tukey, Fisher–Hayter and Wang–Cui methods are designed for use in tests of differences between groups, and do not extend trivially to studies that do not follow an AN(C)OVA design.

After correction for the multiplicity of contrasts, no significant differences were observed between the BSMSS groups. The BSMSS score ranges between 8 and 66 and can be used as a proxy for the social status by assessing, for a child, the occupation of their parents and level of schooling (Barratt, 2006). It does not measure the social class directly, neither the economic status. Although some studies have found brain regions, such as the hippocampus, that appear to be correlated with socioeconomic status (SES), income, and/or stress related to SES (Hanson et al., 2011; Hair et al., 2015; Hanson et al., 2015; Jednoróg et al., 2012; Luby et al., 2013; Yu et al., 2018; Dufford et al., 2018; McDermott et al., 2019), none of them investigated the relation between the brain morphology and only the social status. Even though the score of social status provided by BSMSS and the socioeconomic status are related, they are not equivalent (Barratt, 2006). Besides, other studies classified the subjects as “ in” or “ out” of the poverty class, while in this ANCOVA example we are dividing the subjects into 5 groups using the data as available publicly. Although the first percentile represents the subjects in the lower social classes, this does not imply that they are also in the poverty class. Therefore, the findings without correction shown in Figure 4 might be indeed false positives.

For ECW, more than half of the effects found without correction did not survive after correction using the valid methods (thus, excluding Fisher’s LSD). However, those that did survive, did so consistently across most of the methods, including Dunn–Šidák and Bonferroni. As these are the most conservative (except Scheffé), the fact that the results are generally similar across the methods, with only small differences compared to permutation (Figure 5) and the other valid methods, suggests that it is unlikely that these results are mere false positives. To the best of our knowledge, there are no studies investigating the correlation between the body ECW and the cortical volume of any areas. The possibility that these ECW effects are true positives would be strengthened after correcting for the number of regions considered, that is, the 8 regions from each hemisphere, plus the eTIV (these here take the role of image points). However, doing so in this analysis would unduly punish all methods except Westfall–Young, Bonferroni and Dunn– Šidák, since only the latter three can accommodate directly the MTP-I.

It should be noted that any interpretation of these results, as to whether they represent true positives or not, need to take power into consideration: any approach to correct for multiple testing has the obvious drawback of a drop in statistical power. While one traditionally worries more about type I error, type II error rate is also a consequence of low accuracy of parameter estimates in low powered studies. The noted discrepancies in the literature may be a consequence of random sampling error, since reductions in power are necessary to control the amount of error type I in the multiple testing context. Moreover, presumably much of the published literature on the topic suffers from low statistical power, regardless (for further discussion, see Mumford, 2012; Button et al., 2013; Cremers et al., 2017; Szucs and Ioannidis, 2017).

## 6. Conclusions

We compared different methods for multiple testing correction across contrasts in the context of the GLM, for both synthetic and real data, and argued that such correction is necessary to avoid excess of false positives. Among those methods, Westfall-Young permutation method offers a set of key advantages, some of which were demonstrated. It controls the error rate close to the nominal level, and it is also the most flexible method, as it can be used with arbitrary GLM designs, it corrects over specific hypotheses of interest, and allows correction to various sources of multiplicity, all of which can be implemented in the same algorithm with minimal cost.

## Appendix A. Relation between Dunn–Šidák and Bonferroni

Dunn–Šidák can be seen as a special case from Kimball’s inequality, which states that the probability of all events in a set happening is equal to or greater than the product of their individual probabilities, and where the equality holds only if these events are independent. Dunn–Šidák’s inequality was proved valid for data following a multivariate normal distributions with positive definite correlation matrix (Hochberg and Tamhane, 1987), and therefore, can accommodate some negative correlations between tests. Bonferroni is an approximation of a similar inequality devised by Boole, which establishes the probability of any of the events in a set happening. Although always valid (independently of correlation structure of the tests), it is an approximation and, therefore, is never exact. In general, the relation between Dunn–Šidák and Bonferroni is expressed as (Abdi, 2007)

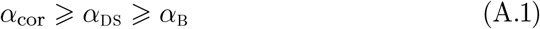

where *α*_DS_ and *α*_B_ are, respectively the Dunn–Šidák and Bonferroni’s corrected test level. The equality between Dunn–Šidák and Bonferroni holds only for *τ* = 1. As more tests are performed, *α*_B_ is always smaller than *α*_DS_, stabilizing at a ratio of approximately 0.975 for an uncorrected test level *α*_unc_ = 0.05, as shown in Figure A.1. The smaller the p-values, smaller is this ratio. If the tests are independent, the correction using Dunn–Šidák is exact, that is, *α*_cor_ = *α*_DS_.

**Figure A.1:**
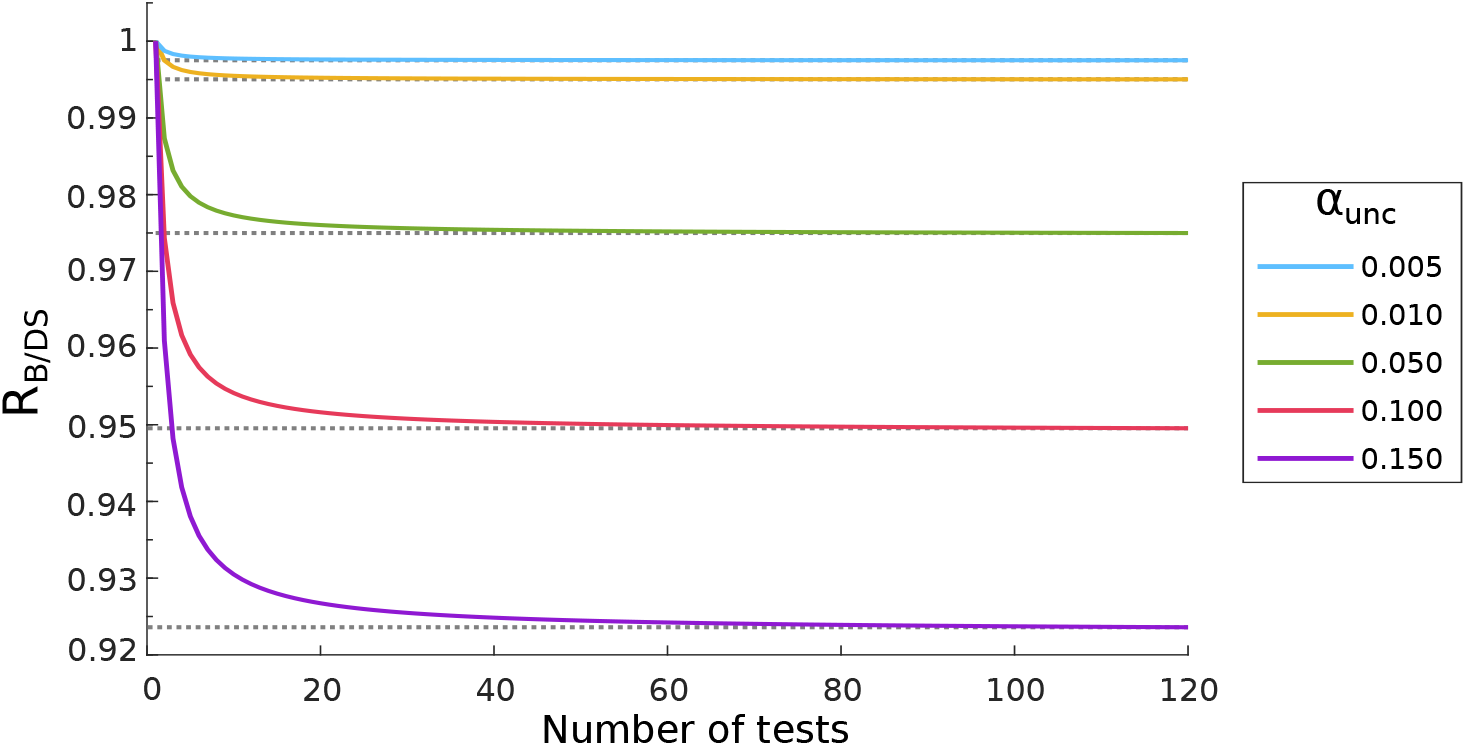
Comparison between the correction performed by Bonferroni and Dunn–Šidák, where R_B/DS_ = *α*_B_/*α*_DS_. Note that the ratio increases with the number of tests and decreases with the uncorrected level *α*_unc_ used.

## Appendix B. Simulated signals with the *t*-statistic

In general, simulated datasets are obtained by sampling random variables that follow a specific distribution (e.g. normal) consistent with the null hypothesis. If the interest is in the alternative hypothesis, a signal with a specific effect size can be added as a function of the desired power *P*, the test statistic, test level and sample size. In the context of the GLM, the sample size is represented in the design efficiency, i.e., effi = (**C**^′^(**X**^′^**X**)^−1^**C**)^1/2^, where **X** and **C** are known, and the effect size is given by *s*_*t*_ = *t*_*s*_(**C**^′^(**X**^′^**X**)^+^**C**)^1/2^, where 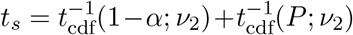 is the location of the peak of signal distribution. Figure B.1 shows two examples of signal added to the normal distribution, one with 50% and the other with 80% power. Note that with 50% power (*P* = 0.50), 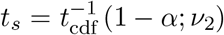 because the peak distribution of the alternative hypothesis is placed over the critical *α* level. In ANOVA designs, where the group means are compared, this signal can be further divided among groups (e.g. half of the signal for each group, as described in Section 3.1).

**Figure B.1:**
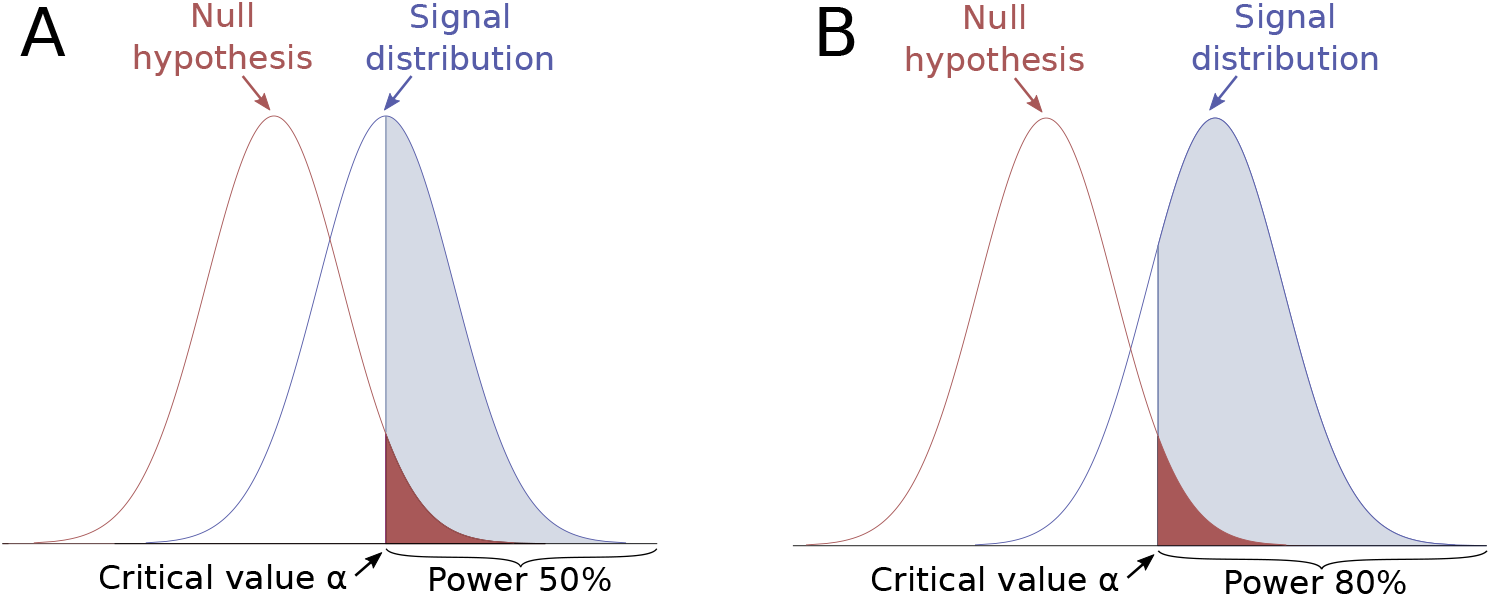
Signal distribution and its relation with the null distribution. Two signals with different powers are exhibited, one with 50% (A) and the other with 80% (B) power.

## Acknowledgements

We would like to thank the anonymous reviewers for their thoughtful comments. We also thank the Child Mind Institute (CMI) for providing data. B.A.V.A. was supported by the Coordenação de Aperfeiçoamento de Pessoal de Nível Superior (CAPES). A.M.W. was supported by the Conselho Nacional de Desenvolvimento Científico e Tecnológico (CNPq; 211534/2013-7). T.E.N. was supported by the Wellcome Trust, 100309/Z/12/Z. This work utilized computational resources of the nih hpc Biowulf cluster (http://hpc.nih.gov).

## Footnotes

1 Both MTP-I and MTP-II refer to the excess of false positives that arises when multiple tests are performed. The MTP-II should not be confused with *type II* errors, which refer to the false negative test results.

2 If multiple image points are also considered, then 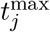 is the maximum value of *t*_*j*_ across both image points and contrasts.

3 For 4, 5, 7 and 9 groups, there are 12, 29, 42 and 72 contrasts being tested. The expected ratio between Bonferroni and Dunn–Šidák is 9.9769 for 12 tests and 9.9751 for 72 tests (see Appendix A).

